# Heparan sulfate molecules mediate synapse formation and function of male mating neural circuits in *C. elegans*

**DOI:** 10.1101/277137

**Authors:** María I. Lázaro-Peña, Carlos A. Díaz-Balzac, Hannes E. Bülow, Scott W. Emmons

## Abstract

The nervous system regulates complex behaviors through a network of neurons interconnected by synapses. How specific synaptic connections are genetically determined is still unclear. Male mating is the most complex behavior in *C. elegans*. It is composed of sequential steps that are governed by more than 3,000 chemical connections. Here we show that heparan sulfates (HS) play a role in the formation and function of the male neural network. Cell-autonomous and non-autonomous 3-*O* sulfation by the HS modification enzyme HST-3.1*/*HS 3-*O*-sulfotransferase, localized to the HSPG glypicans LON-2/glypican and GPN-1/glypican, was specifically required for response to hermaphrodite contact during mating. Loss of 3-*O* sulfation resulted in the presynaptic accumulation of RAB-3, a molecule that localizes to synaptic vesicles, disrupting the formation of synapses in a component of the mating circuits. We also show that neural cell adhesion protein neurexin promotes and neural cell adhesion protein neuroligin inhibits formation of the same set of synapses in a parallel pathway. Thus, neural cell adhesion proteins and extracellular matrix components act together in the formation of synaptic connections.

**Author Summary:** The formation of the nervous system requires the function of several genetically-encoded proteins to form complex networks. Enzymatically-generated modifications of these proteins play a crucial role during this process. These authors analyzed the role of heparan sulfates in the process of synaptogenesis in the male tail of *C. elegans*. A modification of heparan sulfate is required for the formation of specific synapses between neurons by acting cell-autonomously and non-autonomously. Could it be that heparan sulfates and their diverse modifications are a component of the specification factor that neurons use to make such large numbers of connections unique?

## Introduction

Behaviors are the result of a combination of signaling pathways coordinated at various cellular and tissue levels. Male mating is the most complex behavior of *C. elegans*. This behavior is governed by 144 neurons and 64 muscles in the posterior part of the male worm that are extensively inter-connected to each other resulting in approximately 3,200 connected cell pairs (JArrell *et al.* 2012). How these synapses are determined so that they are formed in reproducible patterns is still unknown. One hypothesis is the chemo-affinity hypothesis proposed by Sperry (SPerry 1963), which postulates that matching pairs of cell adhesions molecules between the presynaptic and postsynaptic neurons will promote synaptogenesis. These interactions occur in the context of the surrounding extracellular matrix (ECM), the influence of which needs to be taken into account.

The ECM plays an important role in the development of the nervous system (POrcionatto 2006; ZImmermann And DOurs-ZImmermann 2008; MYers *et al.*2011). Heparan sulfate proteoglycans (HSPG) are components of the ECM that function in several processes such as neurogenesis, cell migration, axon guidance, dendritic branching and synapse formation (BErnfield *et al.* 1999; BÜLow And HObert 2006; POulain And YOst 2015). HSPGs exist in three different forms: (1) transmembrane proteins such as the syndecans; (2) glycosylphosphatidylinositol (GPI) anchored proteins such as the glypicans; and (3) secreted forms such as perlecan, agrin and collagen XVIII (BErnfield *et al.* 1999; BÜLow And HObert 2006; POulain And YOst 2015). A special feature of HSPG is the presence of heparan sulfate (HS) chains that are attached to the core proteins. These HS chains are linear glycosaminoglycan polysaccharides composed of varying numbers of hexuronic acid and glucosamine repeating units (n = 50-150). These polysaccharide chains are subjected to modifications including deacetylation, sulfation, epimerization, and acetylation. The formation of these modifications is catalyzed by heparan sulfate modification enzymes (HSME) (LIndahl And LI 2009). Deacetylation is mediated by N-deacetylase/N-sulfotransferase (Ndst) enzymes, sulfation is mediated by 2-*O*, 6-*O* and 3-*O* HS sulfotransferases, and epimerization is mediated by the glucuronic *C*-5 epimerase. These modifications form domains of distinctly modified HS within the HS chains that serve as binding sites for ligands and receptors. In this way they mediate specific biological functions, such as cell migration and axon guidance (BEnnett *et al.* 1997; PRatt *et al.* 2006; MAtsumoto *et al.* 2007; KAstenhuber *et al.* 2009). Some of these domains may be conserved throughout evolution (ATtreed *et al.* 2016).

The role of HSPG in synapse formation and function is not well-understood. In vertebrates, the HSPG agrin plays a role in the development of neuromuscular junctions by promoting the aggregation of acetylcholine receptors in skeletal muscle and activating the receptor tyrosine kinase Musk on the muscle surface (GLass *et al.* 1996). The presence of agrin has also been detected in the central nervous system, where the suppression of agrin expression in cultured hippocampal neurons and in the cortex of mice results in the formation of fewer synapses (FErreira 1999; BOse *et al.* 2000; KSiazek *et al.* 2007). The intracellular domains of syndecan-2 interact with various cellular components to promote filopodia formation and induce dendritic spine formation (EThell *et al.* 2001; LIn *et al.* 2007). In *Drosophila*, the extracellular and cytoplasmic domains of syndecan act postsynaptically to regulate synapse growth of neuromuscular junctions (NGuyen *et al.* 2016). Glypicans have also been implicated in the formation of synapses. Glypican-4 (Gpc4) and glypican-6 (Gpc6) secretion from astrocytes is sufficient to induce functional synapses in retinal ganglion cells, while their elimination reduces the postsynaptic activity induced by these molecules (ALlen *et al.* 2012). In addition, Gpc4 interaction with a postsynapatic leucine-rich repeat transmembrane protein Lrrtm4 is required to induce excitatory synapse formation (DE WIt *et al.* 2013). This interaction is mediated by an HS-dependent interaction between Gpc4 and the receptor protein tyrosine phosphatase PTPσ in the presynaptic site (KO *et al.* 2015). Additional studies have demonstrated the involvement of HS chain modifications in the process of synapse formation. An RNAi screen in Drosophila, specifically directed to glycan genes, revealed that the functionally paired HS 6-*O* sulfotrasferase (*hs6st*) and HS 6-*O* endosulfatase (*sulf1*) have opposite effects in synaptic functional development of neuromuscular junctions (DAni *et al.* 2012). In mammals, the elimination of *Ext1*, a gene encoding an enzyme essential for heparan sulfate synthesis, causes the attenuation of excitatory synaptic transmission in amygdala pyramidal neurons. In humans, this results in autism-like behavioral deficits (IRie *et al.* 2012).

Here we investigated the role of HS molecules in the development and function of the posterior male nervous system in *C. elegans.* We show that loss of the HS 3-*O* sulfotransferase HST-3.1 and the glypicans LON-2/glypican and GPN-1/glypican result in defects in response to hermaphrodite contact during male mating behavior, suggesting that 3-*O*-sulfated HS attached to LON-2/glypican and GPN-1/glypican is required for this process. In addition, HS molecules and their modifications, with the exception of 3-*O* sulfation, were required for the dorsoventral axonal migration of male-specific sensory neurons that are essential for male mating behavior and function. Loss of 3-*O* sulfation in the postsynaptic cell resulted in accumulation of a presynaptic vesicle marker in the presynaptic cell of a mating circuit. Further, synapse formation between male-specific sensory neurons and target interneurons was disrupted, possibly accounting for the observed behavioral defect.

## Results

### Genetic elimination of HS-modification enzymes and HS proteoglycans causes behavioral defects in male mating

Behavior provides a sensitive readout of developmental or functional disruptions. We conducted a response to hermaphrodite contact behavioral assay of male worms carrying null mutations for the HS-modification enzymes. Response to hermaphrodite contact is the decision of a male worm to start mating after contact by pressing its tail against the hermaphrodite body while moving backward searching for the vulva (Figure 1A). We observed that males carrying a mutation in *hst-2/*HS 2-*O*-sulfotransferase, *hse-5/*HS *C*-5-epimerase and *hst-3.1/*HS 3-*O*-sulfotransferase showed deficiency in response to contact. Loss of *hst-2/*HS 2-*O*-sulfotransferase or *hse-5/*HS *C*-5-epimerase, which respectively introduce 2-*O*-sulfation and *C*-5 epimerization in the hexuronic acid of the linear glycosaminoglycan heparan sulfate chains, mildly affected the response to hermaphrodite contact. Forty five percent of *hst-2(ok595)* and 55% of *hse-5(tm472)* mutant males failed to respond after tail contact compared to a 10% failed response in wild type worms (Figure 1B). For mutants in *hst-3.1(tm734),* which introduces 3-*O*-sulfation, 82% of the *hst-3.1(tm734)* mutant males failed to respond (Figure 1B). This reduction in response to contact results in a reduced number of cross progeny (Figure S1A), demonstrating that the *hst-3.1(tm734)* mutant defect in response reduced the overall ability to succeed in mating. Both response and mating potency defects were rescued by a fosmid containing the *hst-3.1* locus (Figure 1C and S1A). The *hst-3.1(tm734)* mutant deficiencies in response behavior involved a failure in backward movement after mate contact; all non-responsive mutant worms either failed to backup after hermaphrodite contact (52%) or showed a discontinued backward movement losing contact with the hermaphrodite (30%) (Figure S1B). This defect in backward locomotion is not caused by a general defect in backing behavior as *hst-3.1(tm734)* mutant males did not show defects in the backing response to nose touch (Figure S1C). Since *hst-3.1(tm734)* mutant worms showed severe defects in response to contact, subsequent steps of male mating behavior were not examined.

**Figure 1.**
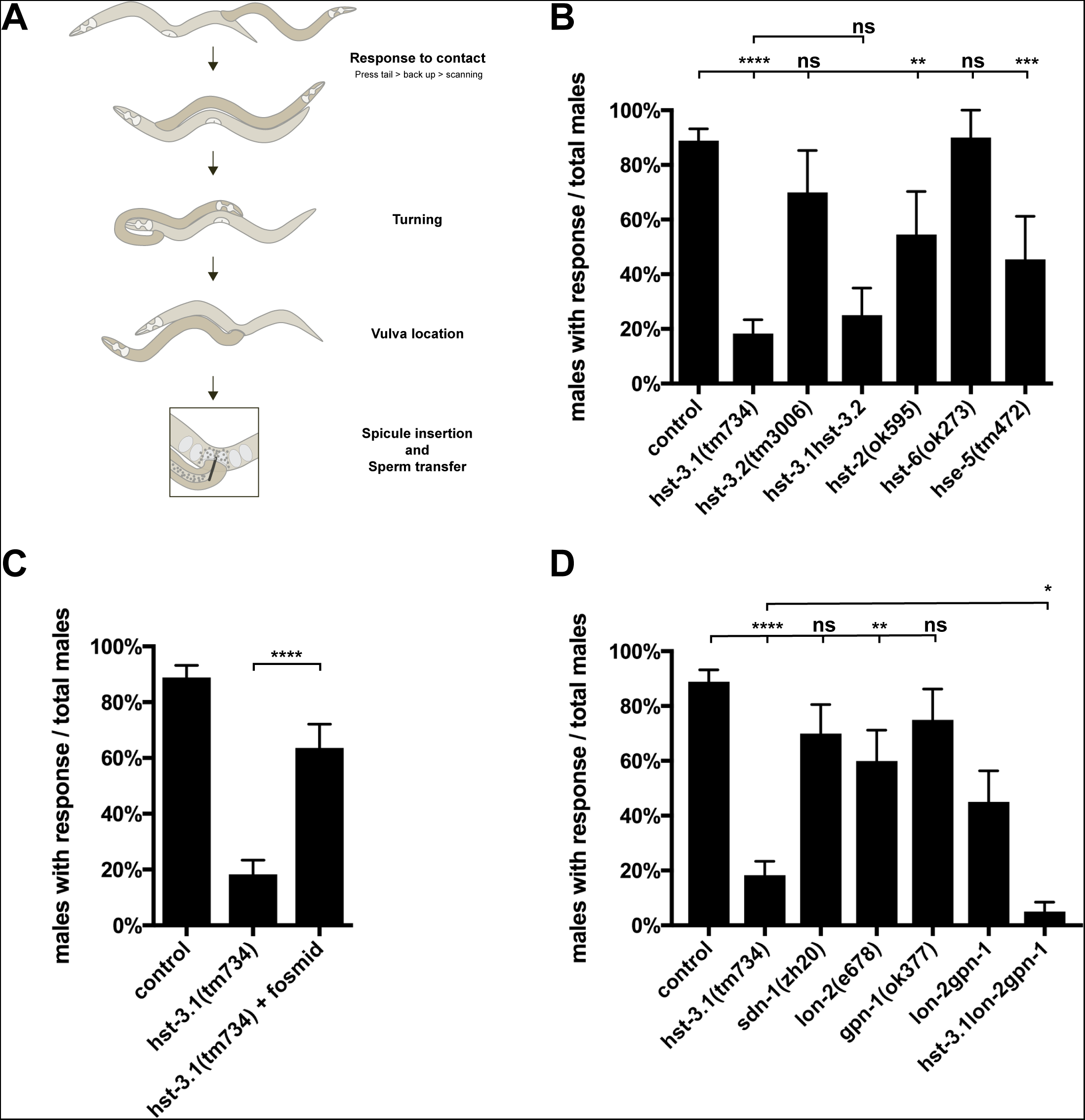
HSME and HSPG are required for response to hermaphrodite contact during male mating behavior. (A) Schematic of the steps of male mating behavior. (B and D) Quantification of response to hermaphrodite contact during male mating behavior in the genotypes indicated. Error bars denote the SEM; statistical significance is shown as follows: *p < 0.05; **p < 0.005; ***p < 0.0005; ****p < 0.00005; and ns, not significant. The data for control are identical (B-D) and shown for comparison only. (C) Quantification of a *hst-3.1*-containing fosmid rescue of the response to hermaphrodite contact during male mating behavior in the *hst-3.1(tm734)* mutant. Error bars denote the SEM; statistical significance is shown as follows: *p < 0.05; **p < 0.005; ***p < 0.0005; ****p < 0.00005; and ns, not significant. The data for control and *hst-3.1* are identical to (B) and shown for comparison only.

The abnormal response to mate contact due to loss of 3-*O*-sulfation was specific for 3-O-sulfation modifications catalyzed by *hst-3.1/*HS 3-*O*-sulfotransferase, since male worms of *hst-3.2(tm3006),* a null allele of the other 3-*O*-sulfatransferase, did not show defects in response after tail contact during mating (Figure 1B). A double mutant for *hst-3.1* and *hst-3.2* did not enhance the abnormal response phenotype of *hst-3.1(tm734)* single mutant, suggesting that 3-*O*-sulfation by *hst-3.1/*HS 3-*O*-sulfotransferase, but not *hst-3.2/*HS 3-*O*-sulfotransferase, is required to mediate male mating behavior. Finally, *hst-6/*HS 6-*O*-sulfotransferase, which introduces 6-*O*-sulfation, is not necessary for response to hermaphrodite contact, as 90% of *hst-6(ok273)* mutant males showed a response after tail contact. Taken together, these results indicate that HS molecules modified by *hst-3.1/*HS 3-*O*-sulfotransferase, *hst-2/*HS 2-*O*-sulfotransferase and *hse-5/*HS *C*-5-epimerase, but not *hst-6/*HS 6-*O*-sulfotransferase, are required for response to hermaphrodite contact by the male worm.

Since *hst-3.1* is a 3-*O*-sulfotransferase that modifies the HS chains on HSPG, we wanted to determine which HSPG may contain the epitope with 3-*O* sulfation that is required for response to hermaphrodite contact. We tested mutants of *sdn-1*/syndecan and the two forms of glypican, *lon-2*/glypican and *gpn-1*/glypican. The *sdn-1(zh20)* mutant males were not defective in response to hermaphrodite contact (Figure 1D). Males containing the *lon-2(e678)* mutation, a hypomorphic allele, were significantly different from control worms as 60% showed a response to contact compared to approximately 90% for control males. This *lon-2* defect is not due to the anatomical Lon phenotype as *lon-1(e185)* mutant males, which have an elongated body similar to *lon-2*, did not show defects in response to hermaphrodite contact (Figure S1D). *gpn-1(ok377)* (a null mutation) mutant males were not significantly different from control worms in their response to hermaphrodites. However, the *lon-2(e678) gpn-1(ok377)* double mutant worms had a stronger mutant phenotype than *lon-2* single mutants, indicating that *gpn-1* also has a function that promotes response (Figure 1D).

To further determine if *hst-3.1/*HS 3-*O*-sulfotransferase is acting on the HS chains attached to *lon-2*/glypican and *gpn-1*/glypican proteoglycans, we constructed triple mutants for *hst-3.1/*HS 3-*O*-sulfotransferase, *lon-2*/glypican and *gpn-1*/glypican. The triple mutant defect was further enhanced as compared to the *hst-3.1/*HS 3-*O*-sulfotransferase single mutant, (Figure 1D), suggesting that LON-2/glypican and GPN-1/glypican have further functions in promoting response independent of 3-*O*-sulfation by *hst-3.1*.

### HS 3-*O* sulfation in the EF interneurons regulates male response to hermaphrodite contact

To define the focus of action of the *hst-3.1/*HS 3-*O*-sulfotransferase HSME in its role in response to contact during mating, we expressed the *hst-3.1/*HS 3-*O*-sulfotransferase cDNA tissue-specifically in neurons, muscles and hypodermis. When we expressed *hst-3.1/*HS 3-*O*-sulfotransferase in neurons using the *rgef-1* pan-neuronal promoter (_*p*_*rgef-1::hst-3.1*) we observed rescue of the abnormal response to contact phenotype (Figure 2A-C). Similarly, when we expressed a *hst-3.1/*HS 3-*O*-sulfotransferase cDNA in hypodermal tissue using the *dpy-7* promoter (_*p*_*dpy-7::hst-3.1*), we observed rescue of the mutant phenotype. However, when we expressed *hst-3.1/*HS 3-*O*-sulfotransferase in muscles using *myo-3* promoter (_*p*_*myo-3::hst-3.1*), we were not able to rescue the response defect. These results suggest that the expression of *hst-3.1/*HS 3-*O*-sulfotransferase in neurons or hypodermis, but not in muscles, is sufficient to regulate response behavior in the male worms.

To further delineate the neuronal focus of action, we used cell-specific promoters that drove expression of the *hst-3.1/*HS 3-*O*-sulfotransferase cDNA in subsets of neurons in the male tale. Cell ablation experiments indicated that the B-type ray sensory neurons are essential for male mating behavior, particularly for the response to hermaphrodite contact and vulva location steps (LIu And STernberg 1995; BArr And STernberg 1999; KOo *et al.* 2011). By examining double mutants, we found that genes such as *pkd-2/*Polycystin-2, *lov-1/*Polycystin-1 and *klp-6*/kinesin that act cell-autonomously in RnB neurons to mediate the response to hermaphrodite contact (BArr *et al.* 2001; PEden And BArr 2005), act genetically in the same genetic pathway as *hst-3.1/*HS 3-*O*-sulfotransferase (Figure S2). However, expression of *hst-3.1/*HS 3-*O*-sulfotransferase in B-type ray neurons by using the *pkd-2/*Polycystin-2 cell-specific promoter (_*p*_*pkd-2::hst-3.1*) was not sufficient to rescue the abnormal response phenotype (Figure 2E). These results suggested that mutation in *hst-3.1* compromised the function of B-type ray neurons but did not act in these neurons. Therefore, to determine whether *hst-3.1/*HS 3-*O*-sulfotransferase is acting downstream of the B-type sensory neurons to regulate response, we expressed the *hst-3.1/*HS 3-*O*-sulfotransferase cDNA in their main postsynaptic partners. Based on the EM male connectivity data, the main postsynaptic targets of RnB neurons are the EF_(1-3)_, PVX, PVY, PVV, PHC and CP_(7-8)_ male-specific interneurons (JArrell *et al.* 2012) (Figure 2D). PVX and PVY are heavily connected to the AVA command interneuron and previous studies revealed that these three interneurons are essential for the backup locomotion that triggers response behavior after mate contact (SHerlekar *et al.* 2013). To express *hst-3.1/*HS 3-*O*-sulfotransferase in the EF_(1-3)_ interneurons, we used *nlg-*1/neuroligin promoter sequence (_*p*_*nlg-1::hst-3.1*), since expression of *nlg-1* in EF_(1-3)_ interneurons has been observed by using a transcriptional GFP fusion (Figure S3C). Expression of *hst-3.1/*HS 3-*O*-sulfotransferase in the EF_(1-3)_ interneurons was sufficient to rescue the abnormal phenotype in response, as approximately 60% of the male worms responded well after tail contact compared to 20% of response in non-transgenic siblings. On the other hand, expression of *hst-3.1/*HS 3-*O*-sulfotransferase cDNA in PVX, PVY, PVV, PHC and AVA interneurons, was not sufficient to rescue the abnormal response phenotype in mating behavior (Figure 2F-I). A previously published transcriptional reporter for *hst-3.1/*HS 3-*O*-sulfotransferase (TEcle *et al.* 2013) is not expressed in EF interneurons (or the hypodermis) and, consistent with this observation, expression of the *hst-3.1/*HS 3-*O*-sulfotransferase cDNA under the same promoter fails to rescue the male mating defects in *hst-3.1/*HS 3-*O*-sulfotransferase mutant males (Figure S3). However, this transgene contains only ∼3kbs of sequences upstream of the start codon and therefore may be missing key regulatory sequences for proper expression of *hst-3.1*. For this reason, we cannot discard the possibility that *hst-3.1/* HS 3-*O*-sulfotransferase might be expressed in the EF_(1-3)_. Altogether, based on the heterologous expression of *hst-3.1/*HS 3-*O*-sulfotransferase, we suggest that *hst-3.1/*HS 3-*O*-sulfotransferase acts in the EF_(1-3)_ male-specific interneurons, which are important post-synaptic targets of B-type ray neurons, to regulate response behavior during male mating.

**Figure 2.**
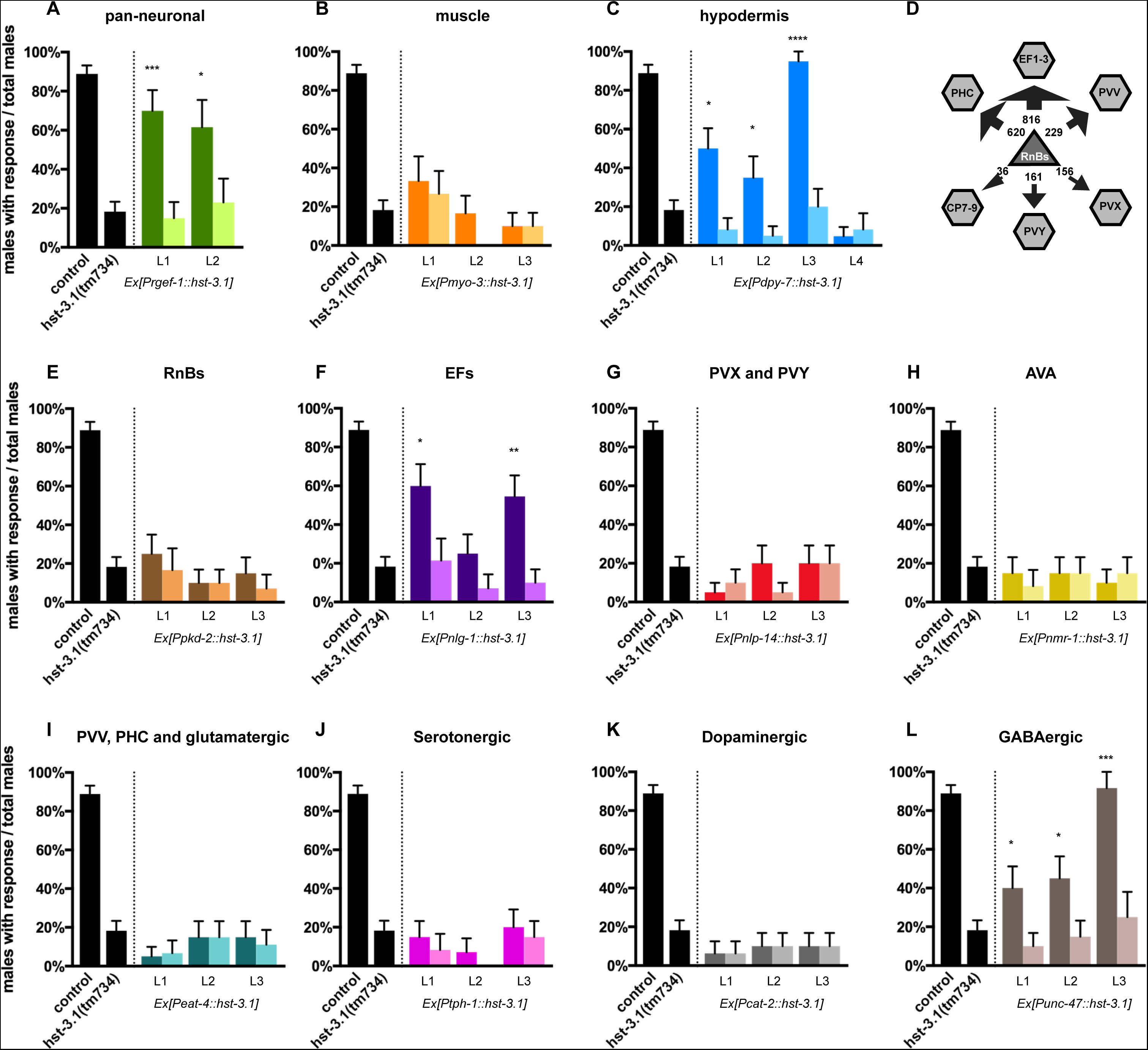
Heterologous Transgenic Rescue Experiments. (A-C) Tissue-specific rescue of response to hermaphrodite contact during male mating behavior in *hst-3.1(tm734)* mutants with *hst-3.1* cDNA under heterologous promoters as indicated. Rescue was defined as restoration of response to hermaphrodite contact during male mating in transgenic animals (darker shade) and had to be statistically significant (p < 0.05) compared to nontransgenic siblings (lighter shade) (n ≥ 12). (D) Main postsynaptic partners of RnB neurons. The arrows and numbers represent the weight of the synaptic input from RnBs to the other neurons. (E-L) Cell-specific rescue of response to hermaphrodite contact during male mating behavior in *hst-3.1(tm734)* mutants with *hst-3.1* cDNA under heterologous promoters as indicated. Rescue was defined as restoration of response to hermaphrodite contact during male mating in transgenic animals (darker shade) and had to be statistically significant (p < 0.05) compared to non-transgenic siblings (lighter shade) (n ≥ 12).

We further tested whether *hst-3.1/*HS 3-*O*-sulfotransferase could act in serotonergic, dopaminergic, glutamatergic and GABAergic neurons by expressing it with the *tph-1* promoter (_*p*_*tph-1::hst-3.1*), *cat-2* promoter (_*p*_*cat-2::hst-3.1*), *eat-4* promoter (_*p*_*eat-4::hst-3.1*) or unc-47 promoter (_*p*_*unc-47::hst-3.1*), respectively. Expression of hst-3.1 cDNA in serotonergic, dopaminergic and glutamatergic neurons did not rescue the abnormal response phenotype (Figure 2I-K). However, expression of *hst-3.1* cDNA in GABAergic neurons rescued the response to hermaphrodite contact defects (Figure 2L). This is consistent with rescue in GABAergic EF interneurons (GEndrel *et al.* 2016).

### HS 3-*O* sulfation is not required for axon guidance of the B-type ray sensory neurons

Since HS molecules are mediators of axon guidance in many neurons in *C. elegans* (BÜLow And HObert 2004; BÜLow *et al.* 2008), we wanted to determine whether the observed behavioral defects in *hst-3.1/*HS 3-*O*-sulfotransferase mutant worms were the result of guidance defects of neurons involved in response behavior. As previously mentioned, the B-type ray sensory neurons are mediators of response to contact behavior and our findings suggest that *hst-3.1/*HS 3-*O*-sulfotransferase is acting from their main postsynaptic target, the EF interneurons, to mediate this behavior. To determine whether 3-*O* sulfation by HST-3.1*/*HS 3-*O*-sulfotransferase in the EF interneurons is acting as a guidance cue to regulate axon migration, we examined B-type ray neurons axonal processes in *hst-3.1/*HS 3-*O*-sulfotransferase mutant worms. To visualize B-type ray neuron processes, we used a cytoplasmic *GFP* reporter driven by the *pkd-2/*Polycystin-2 promoter. *pkd-2/*Polycystin-2 encodes a TRPP cation channel that is expressed in three types of male-specific sensory neurons: the RnBs, HOB and the CEMs.

In wild type worms, during mid to late L4 stage, ray neuron cell bodies migrate away from the posterior tail hypodermis and enter the lumbar ganglion (SUlston *et al.* 1980) (Figure S4A). The axons migrate out of the lumbar ganglion in a DV pathway, forming four to five circumferential commissures. On the ventral side, the axon terminals enter the pre-anal ganglion (PAG) where they contact their postsynaptic partners. In the case of R1B and sometimes R2B, the axons first migrate anteriorly before changing to the DV migration. Therefore, defects in AP migration of B-type ray neurons result in an anterior overextension of processes, while defects in DV migration result in the absence of commissures. As previously reported (JIa And EMmons 2006), these migratory defects are observed in mutant worms for the netrin signaling pathway, where *unc-6(ev400)* and *unc-40(e271)* single mutants showed 30% and 47% (respectively) of defects in AP migration, while both mutants showed 100% of defects in DV migration (Figure 3). We found that HS-modifications 2-*O* sulfation, 6-*O* sulfation and 5-*C* epimerization are required for B-type ray neurons axon guidance as defects are observed in both AP and DV migratory pathways of *hst-2(ok595)*, *hst-6(ok273)* and *hse-5(tm472)* single mutant worms (Figure S4C-D). In addition, SDN-1 is a regulator of B-type ray neurons axon guidance as defects in both AP and DV migration were observed in *sdn-1(zh20)* single mutants. The *hst-2(ok595); hst-6(ok273)* and *sdn-1(zh20); lon-2(e678)* double mutants showed highly penetrant AP and DV migration defects. In fact, the DV migration defects of these double mutants are comparable to those observed in netrin signaling mutants, suggesting that HS molecules and their distinct HS-modification patterns regulate B-type ray neurons axon guidance through the *unc-6*/netrin ligand system (Figure 3 and S4C-F).

**Figure 3.**
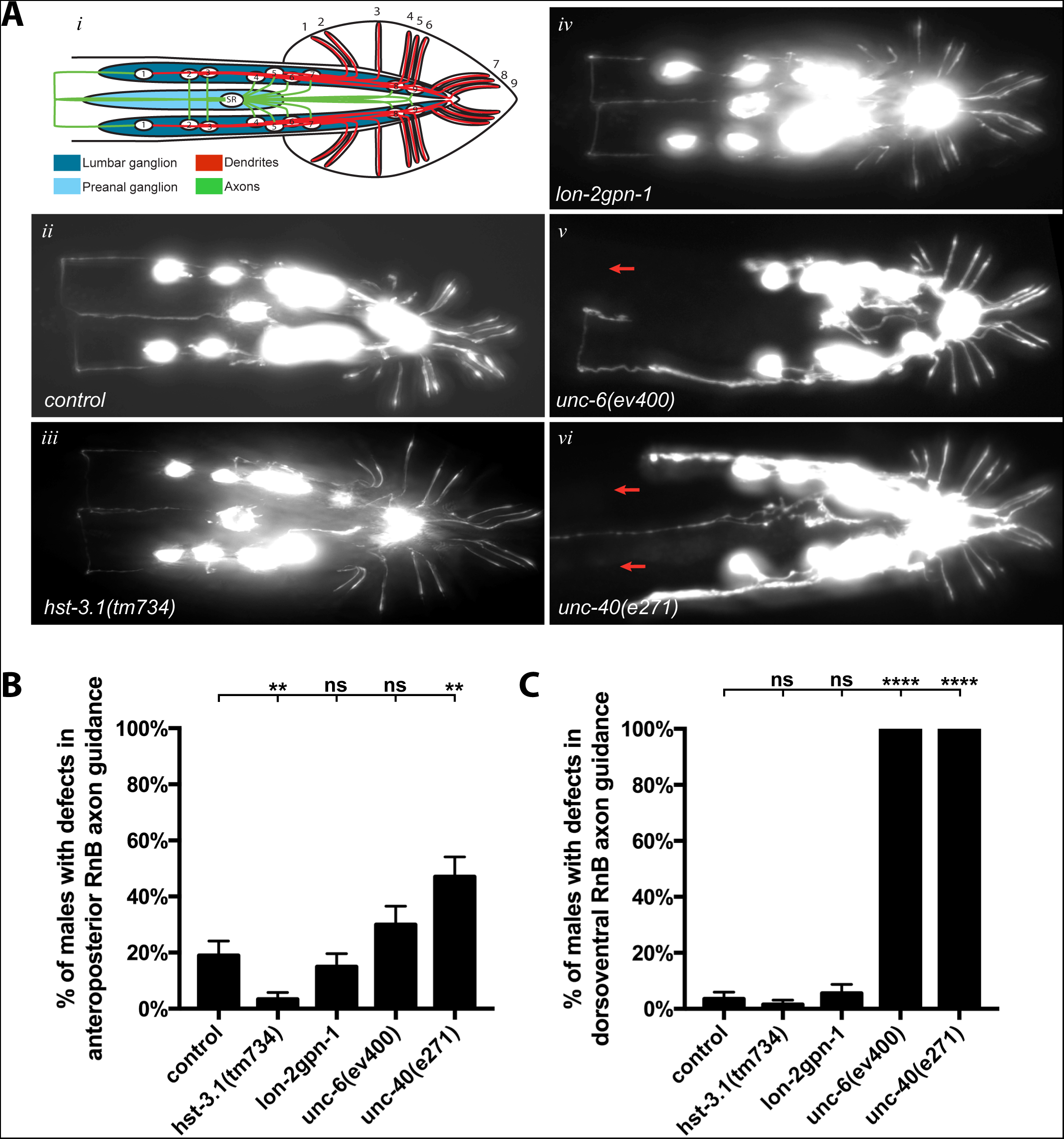
HST-3.1*/*HS 3-*O*-sulfotransferase is not required for axon guidance of B-type ray neurons. (A) Ventral views with schematics (*i-vi*) of adult male animals showing the B-type ray neurons. B-type ray neurons were visualized with *bxIs14* (*Is*[*Ppkd-2::GFP*]). Red arrows represent missing commissures. Anterior is to the left. (B and C) Quantification of B-type ray neurons anteroposterior and dorsoventral axon guidance in the genotypes indicated. Error bars denote the SEM; statistical significance is shown as follows: *p < 0.05; **p < 0.005; ***p < 0.0005; ****p < 0.00005; and ns, not significant.

In contrast, the axon morphology of B-type ray neurons in *hst-3.1(tm734)* single mutant worms was indistinguishable from control worms, with no defects observed in the AP or DV axon migration of RnB neurons (Figure 3). The same results were obtained when we examined the *lon-2 gpn-1* double mutants, which are also defective for response behavior, where no defects in RnB neurons AP or DV axon migration were observed. These results suggest that the defects in response to hermaphrodite behavior in *hst-3.1/*HS 3-*O*-sulfotransferase and *glypican* mutants are not a consequence of defective axonal projections of B-type ray neurons function. Taken together, our analysis of the roles of HS molecules in axon guidance suggests that specific HS-modifications are required for axonal migration of B-type ray neurons, whereas, 3-*O* sulfation by HST-3.1/HS 3-*O*-sulfotransferase is not required for this process, but may rather serve different functions.

### HS 3-*O* sulfation mediates synapse formation of B-type ray sensory neurons

The observation that *hst-3.1/*HS 3-*O*-sulfotransferase mutants are defective in response to contact behavior during male mating, but do not display any obvious defects in axonal projections, prompted us to investigate whether HS 3-*O* sulfation is regulating synaptic function of the response circuits. To examine the possibility that *hst-3.1/*HS 3-*O*-sulfotransferase is affecting synapse formation of B-type ray neurons, we investigated the presynaptic densities of mutant worms using an *mCherry::RAB-3* reporter expressed under the control of *pkd-2/*Polycystin-2 promoter. *rab-3* encodes a member of the Ras GTPase superfamily that localizes to presynaptic sites. To study RnB synapses, we used the confocal microscope and analyzed the synaptic pattern of compressed z-stacks containing all the RnBs synapses located in the preanal ganglion. We looked at the synapses of young adult male worms (64 hours after hatching) because by this time most B-type ray neuron synapses are formed (Figure S5A). Based on the presynaptic puncta distribution in the preanal ganglion, the presynaptic density patterning of *hst-3.1/*HS 3-*O*-sulfotransferase mutant males was similar in morphology to that of control worms, suggesting that the elimination of *hst-3.1/*HS 3-*O*-sulfotransferase does not affect the distribution of B-type ray neurons presynaptic sites (Figure 4B).

**Figure 4.**
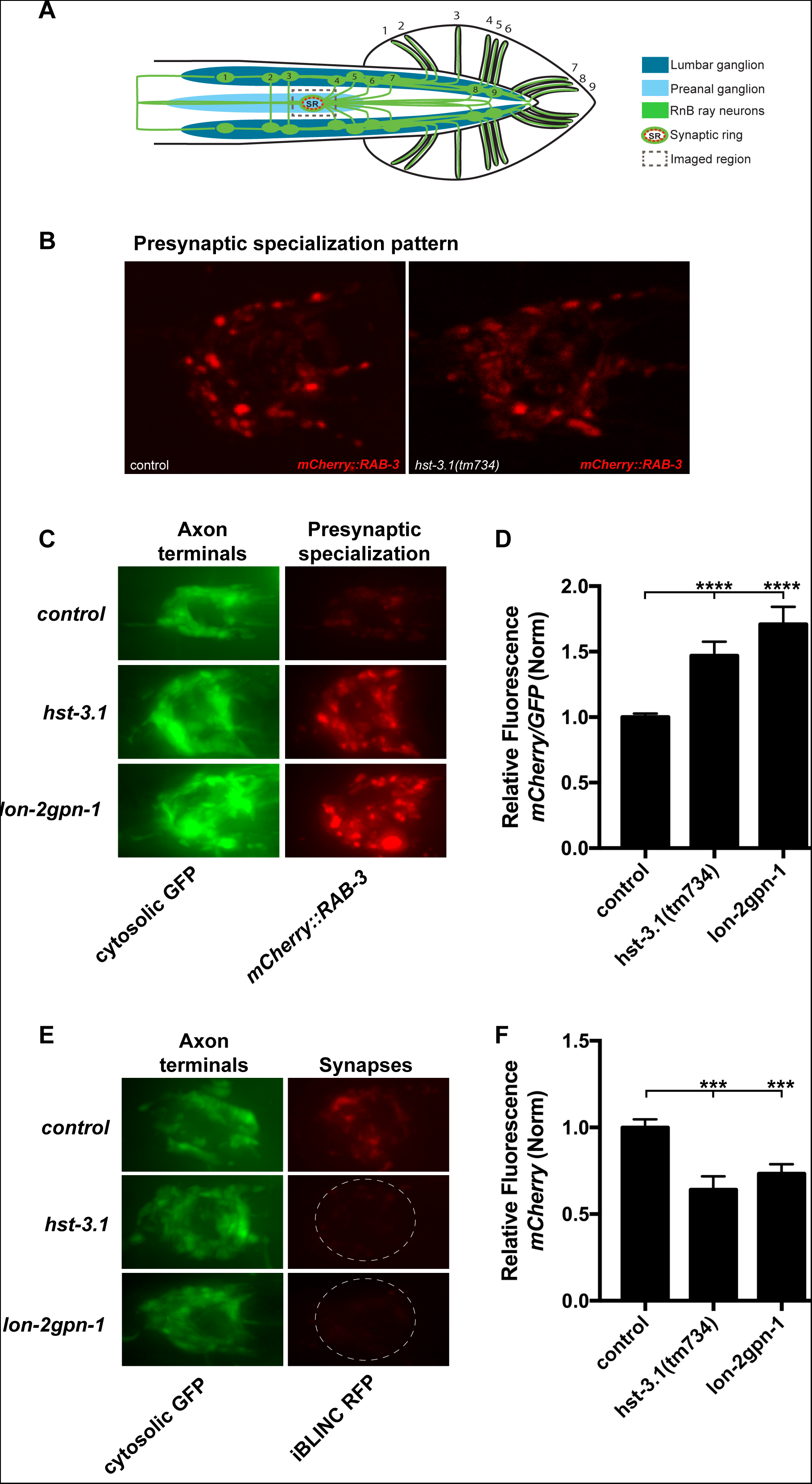
HST-3.1*/*HS 3-*O*-sulfotransferase regulates presynaptic organization and synapse formation of B-type ray neurons. (A) Schematic of a ventral view showing the imaged region containing the synaptic ring in the preanal ganglion. (B) Confocal ventral views of the presynaptic densities as labeled with mCherry::RAB-3 of adult male animals in control and *hst-3.1* mutant. The B-type ray neurons presynaptic distribution is not affected in *hst-3.1/*HS 3-*O*-sulfotransferase mutants. (C) Ventral views of the RnB axonal processes in the synaptic ring located in the preanal ganglion with cytosolic GFP (A-C) and its corresponding presynaptic densities as labeled with mCherry::RAB-3 (A′-C′) of adult *hst-3.1(tm734)* single mutants and *lon-2gpn-1* double mutant male worms. B-type ray neurons were visualized with *bxIs30* that contains the cytosolic GFP (*Is*[*Ppkd-2::GFP*]) and the presynaptic marker (*Is*[*Ppkd-2::mCherry::RAB-3*]). Anterior is to the left. (D) Quantification of mCherry::RAB-3 fluorescence in the preanal ganglion synaptic ring in the genotypes indicated. Error bars denote the SEM; statistical significance is shown as follows: *p < 0.05; **p < 0.005; ***p < 0.0005; ****p < 0.00005; and ns, not significant. The data presented is a ratio of mCherry::RAB-3 to GFP and control value. (E) Ventral views of trans-synaptic biotinilation labeling (iBLINC) of RnB > EFs synapses in puncta in *hst-3.1(tm734)* single mutants and *lon-2gpn-1* double mutants. (F) The data presented is the normalized RFP density in the synaptic area. Error bars denote the SEM; statistical significance is shown as follows: *p < 0.05; **p < 0.005; ***p < 0.0005; ****p < 0.00005; and ns, not significant.

However, we found that the *mCherry::RAB-3* levels are accumulated above control levels (Figure 4C). To quantify the levels of *mCherry::RAB-3* in synapses we performed a densitometry analysis of compressed z-stack images containing all RnB synapses in the preanal ganglion. To control for variability in transgene expression, we normalized by using a cytoplasmic GFP expressed from the same transgene and promoter. We observed that *hst-3.1/*HS 3-*O*-sulfotransferase mutants have a *mCherry* relative fluorescence of 1.47 ± 0.11 compared to 1 ± 0.03 in control worms (Figure 4D).

The *mCherry::RAB-3* levels in the B-type neuron synapses of *lon-2 gpn-1* double mutant worms have a similar accumulation of RAB-3 with a relative fluorescence of 1.71 ± 0.13 compared to 1 ± 0.03 in control worms (Figure 4D). The *lon-2(e678)* single mutant worms, which are defective for response to contact, also showed increased levels of RAB-3 in B-type ray neuron synapses when compared to control (Figure 1D and S6B).

Interestingly, mutant worms for the other HSME with defects in male mating behavior, *hse-5(tm472)*, also showed higher *mCherry::RAB-3* accumulation levels significantly different from control worms (Figure 1B and S6A). However, given that these mutants are also defective in axon guidance (Figure S4C-D), the observed behavioral defects might be due to axon misrouting rather than synaptic function, as seems to be the case for *hst-3.1/*HS 3-*O*-sulfotransferase. Finally, *hst-3.2(tm3006)* and *hst-6(ok273)* mutant worms did not show defects in the *mCherry::RAB-3* accumulation in synapses, nor did they show defect in response to contact behavior as shown above (Figure S6A and 1B). The single and double HSPG mutants such as *sdn-1(zh20), gpn-1(ok377)*, *unc-52(e998)*, *sdn-1 lon-2*, and *sdn-1 gpn-1*, did not show different levels of accumulation of *mCherry::RAB-3* in presynaptic sites compared to control worms (Figure S6B).

The abnormal *mCherry::RAB-3* accumulation observed in B-type ray neurons of *hst-3.1/*HS 3-*O*-sulfotransferase and *lon-2 gpn-1* mutant worms suggests a synaptic disruption between these neurons and their postsynaptic partners. To examine this possibility, we used the iBLINC (Biotin Labeling of INtracellular Contacts) system to label the specific synapses between B-type ray neurons and EF interneurons. The iBLINC method consists of an enzymatic trans-synaptic transfer reaction of biotin from a presynaptic cell adhesion protein to a postsynaptic molecule (DEsbois *et al.* 2015). The biotinylated postsynaptic site is detected by streptavidin::fluorescent protein, thereby labeling the synapse (Figure S5B). For the B-type ray neurons, we expressed a BirA ligase fused N-terminally to nrx-1/neurexin driven by the *pkd-2/*Polycystin-2 promoter. For the EF interneurons, we expressed an acceptor peptide AP fused N-terminally to nlg-1/neuroligin driven by the *nlg-1*/neuroligin promoter. We found that *hst-3.1(tm734)* mutants show less synaptic labeling compared to control worms, suggesting smaller or fewer synapses between RnBs and the EFs neurons. Control worms have a relative fluorescence of 1 ± 0.05, while the *hst-3.1(tm734)* mutants have a relative fluoresce of 0.26 ± 0.03 (Figure 4E and 4F). Moreover, *lon-2 gpn-1* double mutant worms also showed less synaptic labeling than control worms with a relative fluorescence of 0.75 ± 0.05. Altogether, our synaptic and behavioral results suggest that HS 3-*O* sulfation is involved in the process of synapse formation, which in turn may affect the function of the connection between B-type ray neurons and EF interneurons, resulting in the defects in response behavior during male mating.

### HST-3.1*/*HS 3-*O*-sulfotransferase and the glypicans, LON-2/glypican and GPN-1/glypican, genetically interact with synaptic molecules

The process of synapse formation is thought to be mediated by the interaction of cell adhesion molecules located in the presynaptic and postsynaptic sites. Such is the case for presynaptic neurexin and postsynaptic neuroligin adhesion molecules, which have been implicated in synaptogenic activity and synapse maturation (SCheiffele *et al.* 2000; GRaf *et al.* 2004; SÜDhof 2008). We found *nrx-1*/neurexin null mutants exhibited defects in response to contact during male mating similar to the defects observed in *hst-3.1/*HS 3-*O*-sulfotransferase mutant worms (Figure 5A). *nlg-1*/neuroligin mutants, by contrast, did not show defects in response to contact behavior. To investigate whether *hst-3.1* still retained a function in the absence of *nrx-1* function, we constructed *hst-3.1; nrx-1* double mutants. We found that the *hst-3.1; nrx-1* double mutant further enhanced the defects observed in both single mutants, thus suggesting that these genes act in parallel genetic pathways, while not excluding the possibility that they act in the same pathway given the strong phenotype observed in the *hst-3.1/*HS 3-*O*-sulfotransferase single mutant (Figure 5A).

**Figure 5.**
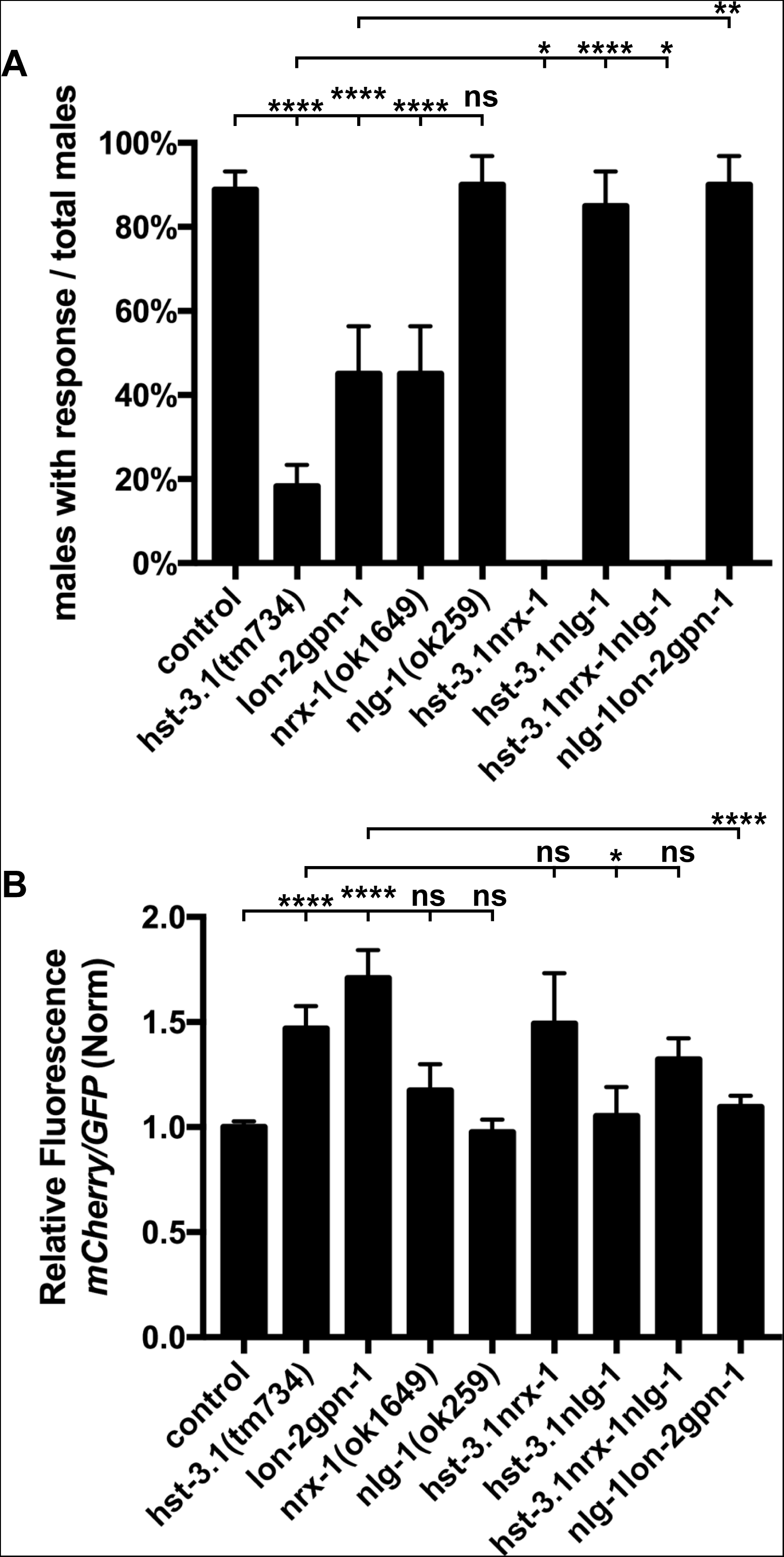
*nrx-1*/neurexin and *nlg-1*/neuroligin interacts genetically with *hst-3.1/*HS 3-*O*-sulfotransferase for response to hermaphrodite contact and synaptic function. A) Quantification of response to hermaphrodite contact during male mating behavior in the genotypes indicated. Error bars denote the SEM; statistical significance is shown as follows: *p < 0.05; **p < 0.005; ***p < 0.0005; ****p < 0.00005; and ns, not significant. The data for control and *hst-3.1/* HS 3-*O*-sulfotransferase are identical to figure 1 and shown for comparison only. B) Quantification of mCherry::RAB-3 fluorescence in the preanal ganglion synaptic ring in the genotypes indicated. The data presented is a ratio of mCherry::RAB-3 to GFP and control value. Error bars denote the SEM; statistical significance is shown as follows: *p < 0.05; **p < 0.005; ***p < 0.0005; ****p < 0.00005; and ns, not significant. The data for control and *hst-3.1/* HS 3-*O*-sulfotransferase are identical to figure 4 and shown for comparison only.

The *hst-3.1; nlg-1* double mutant suppressed the defect in response to contact during male mating observed in the *hst-3.1* single mutant (Figure 5A). Thus there is an *hst-3.1*-independent pathway that is suppressed by *nlg-1*. This pathway depends on *nrx-1* function. We examined *hst-3.1; nrx-1; nlg-1* triple mutant worms and observed that in the absence of *nrx-1*/neurexin function, a mutation in *nlg-1*/neuroligin no longer suppresses the response defects due to mutation in *hst-3.1*. Thus *nrx-1*/neurexin is epistatic and suggesting that *nlg-1* acts to suppress the activity of *nrx-1* that promotes response behavior. To further investigate the *nlg-1*/neuroligin suppression of defects in response behavior due to mutation in *hst-3.1*, we constructed triple mutants for *nlg-1*/neuroligin and the two glypicans, *lon-2*/glypican and *gpn-1*/glypican. The *nlg-1 lon-2 gpn-1* triple mutants did not show defects in response to contact, thus indicating that *nlg-1*/neuroligin also suppresses the defects of *lon-2 gpn-1* double mutants (Figure 5A). Together, these results suggest that *nrx-1*/neurexin and *nlg-1*/neuroligin adhesion, possibly through opposing roles, are involved in regulating response behavior in a pathway parallel to that in which *hst-3.1/*HS 3-*O*-sulfotransferase and the glypicans act.

To determine whether the genetic interaction of these molecules is also regulating synapse formation, we examined the presynaptic levels of *mCherry::RAB-3* in B-type ray sensory neurons in the same series of *nrx-1*/neurexin and *nlg-1*/neuroligin double and triple mutant worms. The *hst-3.1; nrx-1* mutants showed an increased accumulation of *mCherry::RAB-3*, however the increase in accumulation was not significantly different from *hst-3.1/*HS 3-*O*-sulfotransferase single mutant worms (Figure 5B). Considering that *hst-3.1; nrx-1* double mutants’ defects in behavior are significantly more severe than the defects in *hst-3.1/*HS 3-*O*-sulfotransferase single mutants, we conclude that either it is difficult to detect an even higher accumulation of *mCherry::RAB-3* as measured by the relative fluorescence densities, or that behavioral defects due to *nrx-*1 mutation result from a disruption in another synaptic connection.

Consistent with the behavioral results, the *hst-3.1; nlg-1* double mutant suppressed the *mCherry::RAB-3* accumulation in the presynaptic cell, decreasing it to control levels (Figure 5B). As we observed in response behavior, the *hst-3.1; nrx-1; nlg-1* triple mutants showed an increased accumulation of *mCherry::RAB-3* that was not suppressed by *nlg-1*/neuroligin, consistently indicating that in the absence of *nrx-1*, *nlg-1* does not suppress the *hst-3.1/*HS 3-*O*-sulfotransferase induced defects in B-type ray neuron presynaptic sites. Lastly, *nlg-1 lon-2 gpn-1* triple mutants showed *mCherry::RAB-3* levels comparable to control worms, again mirroring the genetic interactions observed for the male mating behavior.

## Discussion

How neuronal connectivity determines behavioral output in an organism remains one of the biggest questions in the neuroscience field. To address this question, one important aspect is to investigate the roles of molecules in the extracellular matrix that are involved in establishing and making these connections functional. In this work, we used male mating behavior in *C. elegans* as a read-out of synaptic function together with fluorescence labeling of synapses to study the role of HS molecules in formation of the male nervous system and its synaptic connectivity.

### HS molecules mediate male mating behavior in *C. elegans*

We found that *hst-3.1/*HS 3-*O*-sulfotransferase is acting in the same genetic pathway as *pkd-2/*Polycystin-2, *lov-1/*Polycystin-1 and *klp-6/*Kinesin to regulate response to contact behavior during male mating, implicating compromise of B-type ray sensory neuron function as the basis of the phenotype. However, *hst-3.1/*HS 3-*O*-sulfotransferase focus of action is not the B-type ray neurons, but rather the downstream EFs male-specific interneurons or the hypodermis. Based on the double mutant defects, the HS proteoglycans LON-2/glypican and GPN-1/glypican act in parallel to mediate response behavior. Since it has been shown that *gpn-1*/glypican is expressed in neurons (HUdson *et al.* 2006) and *lon-2*/glypican acts in the hypodermis to mediate different aspects of neuronal development (PEdersen *et al.* 2013), we propose that response to contact behavior is mediated by 3-*O* sulfation to the HS attached to GPN-1/glypican in neurons, and 3-*O* sulfation to the HS attached to LON-2/glypican in hypodermis. Even though glypicans possess a GPI anchor, it has been shown that LON-2/glypican can be shed from epidermal cells, secreted and diffuse into the extracellular matrix where they interact with components of Netrin signaling to mediate axon guidance (BLanchette *et al.* 2015).

Interestingly, only 3-*O*-sulfation by *hst-3.1/*HS 3-*O*-sulfotransferase and not by *hst-3.2/*HS 3-*O*-sulfotransferase is regulating the mating process. The opposite specificity for the 3-*O* sulfotransferases was identified in the neurite branching induced due to overexpression of the cell adhesion molecule KAL-1/Anosmin 1 in the AIY interneuron (BUlow *et al.* 2002). In this context, the *kal-1*-induced branches in AIY were suppressed in the *hst-3.2* mutant, but not in the *hst-3.1/*HS 3-*O*-sulfotransferase mutant (TEcle *et al.* 2013), thus providing further evidence that both HS 3-*O-*sulfotransferase might display different substrate specificities or distinct expression patterns (MOon *et al.* 2012).

It has been shown that the EFs interneurons are important for male exploratory behavior, which is essential for males to locate and contact their mating partners (BArrios *et al.* 2008). In terms of connectivity, their main synaptic input is from the B-type ray sensory neurons, while their main synaptic output is onto the AVB pre-motor interneuron (S.J. Cook, C.A. Brittin, T.A. Jarrell, Y. Wang, A.E. Bloniarz, et al., manuscript in preparation). Previous cell ablation studies have shown that the Efs interneurons mediate backward locomotion after mate contact (SHerlekar 2015), which is necessary for response to contact behavior. Because AVB is the pre-motor interneuron that promotes forward movement in the locomotion circuit, while EF ablation promotes forward movement after mate contact, it is thought that the EFs -> AVB connections are inhibitory synapses. Our findings support this hypothesis given that *hst-3.1/*HS 3-*O*-sulfotransferase is acting in the EFs to regulate response behavior, and the *hst-3.1/*HS 3-*O*-sulfotransferase mutants have defects in backward locomotion after contacting the hermaphrodite.

### HS 3-*O* sulfation regulates synapse formation of male mating neurons

Our findings, together with previously published studies, demonstrate that heparan sulfates are mediators of synapse development and function. In this work, we identify a HS motif with 3-*O*-sulfation likely attached to the HSPG LON-2/glypican and/or GPN-1/glypicans that is required for synaptogenesis. Our observations conceptually extend studies in cultured hippocampal neurons, where the LRRTM4’s synaptogenic activity requires the presence of heparan sulfates (de Wit et al., 2013). Moreover, they showed that glypican acts as a receptor for LRRTM4, and their interaction is important for the development of excitatory synapses.

Through the analysis of null mutations, we demonstrate that the elimination of *hst-3.1/*HS 3-*O*-sulfotransferase and the two glypican forms, *lon-2*/glypican and *gpn-1*/glypican, induces defects in the presynaptic specialization of B-type ray neurons, while it reduces the iBLINC synaptic labeling of RnBs -> EFs synapses, indicating their role in the process of synapse formation. Higher levels of the presynaptic marker RAB-3 in B-type ray neurons in *hst-3.1/*HS 3-*O*-sulfotransferase mutants could be because vesicle fusion is not occurring properly at the synapse. The higher levels of RAB-3 can be explained by a change in the size and morphology of the synaptic puncta rather than an increase in their number. This is suggested by the fact that the presynaptic pattern of B-type ray neurons in the pre-anal ganglion of mutant worms is comparable to the one observed in control worms. In addition, using the iBLINC labeling system, we showed that *hst-3.1/*HS 3-*O*-sulfotransferase mutants and *lon-2 gpn-1* double mutants form fewer RnB -> EFs connections. In terms of synaptic function, from our results, we argue that the observed *hst-3.1/*HS 3-*O*-sulfotransferase and *lon-2gpn-1* defect in response behavior during mating is a reflection of the observed defects in synapse formation.

The HS 3-*O* sulfation regulation during neuronal development of B-type ray sensory neurons seems to be specific for the process of synapse formation as no defects were observed in the axon morphology of these mating neurons. A similar restricted role of HS molecules in the process of synapse formation has been previously reported in mammals, where ext1 conditional knockout in mice results in autism-like behavioral phenotypes due to abnormal functioning of glutamatergic synapses while no detectable morphological defects were observed in the brain (IRie *et al.* 2012). Even though there is no involvement of HS 3-*O* sulfation in the process of B-type ray neurons axon guidance, we found that other HS modifications such as 2-*O* sulfation, 6-*O* sulfation and *C*-5 epimerization mediate anteroposterior and dorsoventral axon guidance pathways by acting in parallel genetic pathways. This is consistent with previous findings that demonstrate the various functions of distinct HS modification patterns in the development of different neuronal cell types (SAied-Santiago *et al.* 2017). The simultaneous knockdown of three proteoglycans, *sdn-1*/syndecan, *lon-2*/glypican and *gpn-1*/glypican, severely affected axon guidance of B-type ray neurons demonstrating they act redundantly in this process, as is the case for other processes such as KAL-1/Anosmin 1 induced neurite branching in the AIY interneuron (DIaz-Balzac *et al.* 2014). In the context of dorsoventral migration of B-type neurons, genetic elimination of HS molecules causes defects similar to *unc-6*/netrin ligand and its *unc-40*/DCC surface receptor; possibly HS may regulate axon guidance through the netrin signaling pathway. However, the mechanism by which this is accomplished remains elusive. Plausible possibilities include the ligand sequestration, the modulation of ligand-receptor interaction, or a function of HSPGs as co-receptors (BLanchette *et al.* 2015; DIaz-BAlzac *et al.* 2015; POulain And YOst 2015).

### A role for neurexin and neuroligin

We have shown that neural cell adhesion proteins neurexin and neuroligin play a role in the formation of synapses by the RnB neurons. Neurexin appears to promote synapse formation while neuroligin opposes the function of neurexin. Interestingly, Hart and Hobert (HArt And HObert 2018) recently found a similar novel, mutually antagonistic role for these proteins in synapse formation elsewhere in the *C. elegans* male mating circuits. This neurexin/neuroligin pathway is to some extent independent of the pathway in which *hst-3.1/* HS 3-*O*-sulfotransferase functions as each gene retains some function in null mutants of the other (Figure 6). In fact, in the absence of the inhibitory function of neuroligin, it appears that the synapse-promoting function of neurexin can fully restore synapse formation and function in the absence of *hst-3.1/* HS 3-*O*-sulfotransferase (Figure 5). These results indicating multiple independent pathways promoting synapse formation point up the complexity of the process and help to understand how specificity and robustness are achieved.

**Figure 6.**
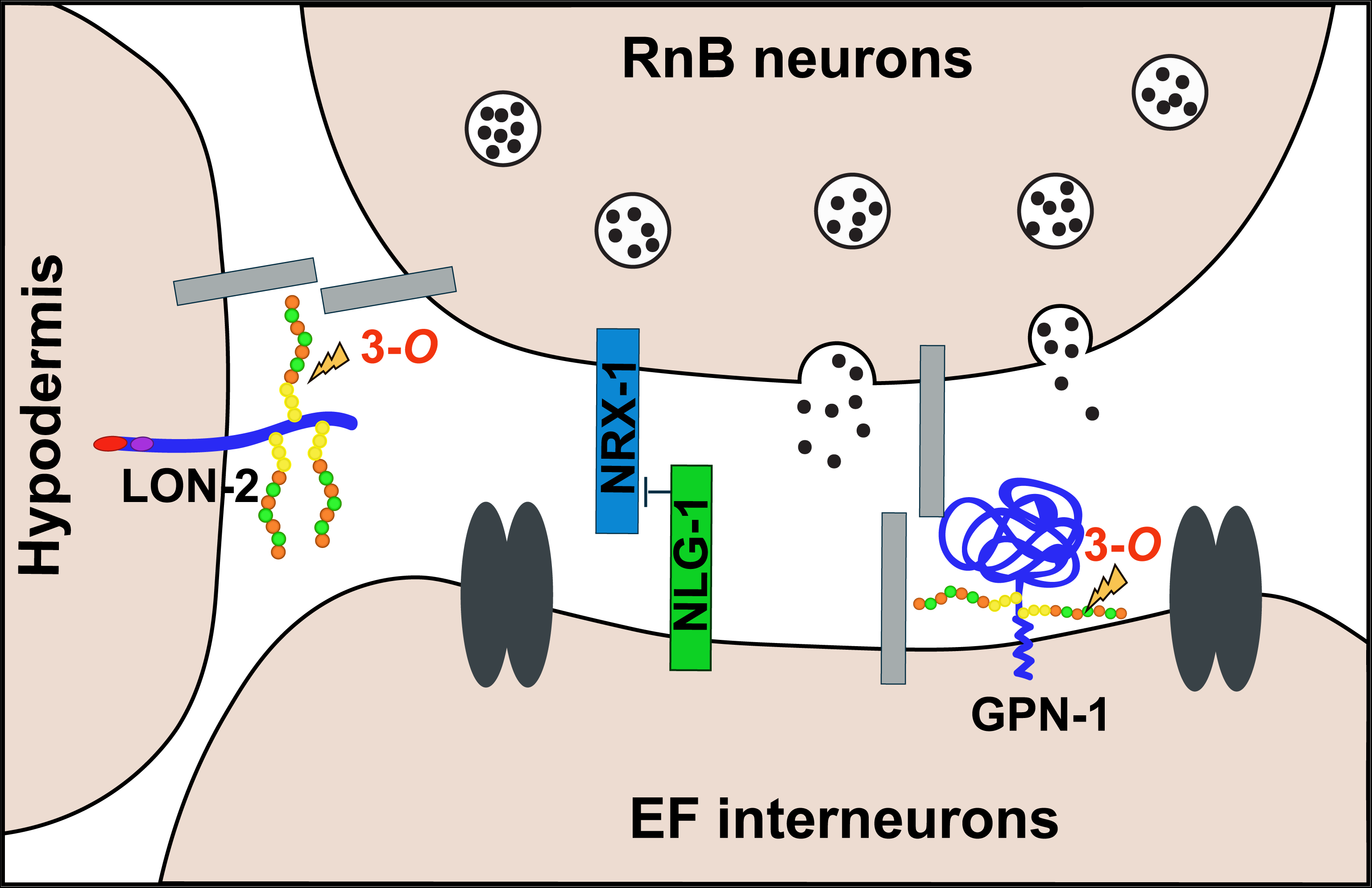
Proposed working model for the role of HS 3-*O* sulfation in synapse formation. 3-*O* sulfation of HS chains located in LON-2 and GPN-1 mediates synapse formation between RnB neurons and EF interneurons, in parallel to NRX-1 and NLG-1, most likely by regulating the interaction between unidentified synaptic molecules. The absence of NRX-1 promotes synapse formation between RnB neurons and EF interneurons, while NLG-1 acts as a synaptic inhibitor. Disruption of this synaptic connection causes the accumulation of synaptic vesicles in the B-type ray neurons and behavioral defects in response to hermaphrodite contact during male mating. However, further biochemical experiments are needed to validate this model.

## Acknowledgements

We thank the laboratory members of the Emmons and BÜlow lab for comments on the manuscript and helpful discussions; the Caenorhabditis Genetics Center (which is funded by the NIH, P40 OD010440) for strains; Oliver Hobert Lab for DNA clones. This work was supported in part by grants from the National Institutes of Health (T32 GM007491 to M.I.L.P.; T32 GM007288 and F31 HD066967 to C.A.D.B.; RC1 GM090825 and R01 GM101313 to H.E.B.; and R01 GM066897 to S.W.E.) and the G. Harold & Leila Y. Mathers Charitable Foundation (to S.W.E). H.E.B. is an Irma T. Hirschl/Monique Weill-Caulier Research Fellow.

## Experimental Procedures

### *C. elegans* strains and imaging

All strains were maintained using standard methods (BRenner 1974). All strains used contain the *him-5(e1490)* mutation on chromosome V to increase the male population (Broverman 1994). We refer to *him-5* as control worms. All experiments were performed at 20°C, and animals were scored as 1-day-old adults unless otherwise specified. The strains and mutant alleles used in this study are listed in the Supplemental Experimental Procedures. Fluorescent images were captured in live *C. elegans* using a Plan-Apochromat 40×/1.4 or 63x/1.4 objective on a Zeiss Axioimager Z1 Apotome. Worms were immobilized using 10 mM sodium azide and *Z* stacks were collected. Maximum intensity projections were used for further analysis.

### Molecular biology and transgenesis

To assemble tissue specific expression constructs used for rescue experiments, the *hst-3.1* cDNA was cloned under control of the following promoters: hypodermal *Pdpy-7* (GIlleard *et al.* 1997), body wall muscle *Pmyo-3* (OKkema *et al.* 1993), pan-neuronal *Prgef-1* (ALtun-Gultekin et al. 2001), the B-type ray neurons *Ppkd-2 (BArr And STernberg 1999)*, EF interneurons *Pnlg-1* (this study), dopaminergic neurons *Pcat-2* (LInts And EMmons 1999), PVY and PVX neurons *Pnlp-14* (SHerlekar *et al.* 2013) AVA neuron *Pnmr-1* (SHerlekar *et al.* 2013) glutamatergic neurons *Peat-4* (LEe *et al.* 1999), serotonergic neurons *Ptph-1* (SZe *et al.* 2000), and gabaergic neurons *Punc-47* (GEndrel *et al.* 2016). All plasmids contained the *unc-54* 3’UTR. Constructs for tissue specific rescue experiments of *hst-3.1/* HS 3-*O*-sulfotransferase male mating response defect were injected at 5 ng/µl together with *Pceh-22::GFP* or *Punc-122::GFP* as injection markers at 50 ng/µl. For details see Supplemental Experimental Procedures.

### Behavioral scoring

Response to hermaphrodite contact assays were performed with one day old adult, virgin males isolated at the L4 stage. Male worms to be tested were placed in a 10-mm food lawn with ten *unc-31(e169)* hermaphrodites. The mating behavior of the males was observed for 5 minutes and annotations were made every time a male responded to hermaphrodite contact. For this assay, a male is considered to have responded to contact if after mate contact it starts backward locomotion, scanning the hermaphrodite body followed by turning behavior. Males that failed to response to contact did not start backward locomotion to scan the hermaphrodite body, or they lost tail contact right after starting the backward locomotion. The quantitation of response to contact was performed by dividing the number of males with response by the total number of male tested [Response % = (# of males with response/total # of males evaluated) x 100. For statistical analysis, we performed a t-test to calculate the significant difference between control worms (*him-5*) and mutant worms (in a *him-5* background).

For the mating potency assay, we placed one virgin young adult male worm with one *pha-1* mutant hermaphrodite in a 10-mm food lawn for 4 hours. We scored a total of 50 male worms (50 plates). After 4 hours, males were removed from the plate and the plate was placed at 25°C. The *pha-1* mutant worms are temperature sensitive and are not viable at 25°C, so only the crossed progeny grows at 25°C. Three days later, we counted the plates with worms that survived at 25°C. To calculate mating potency, we divided the number of cross progeny plates by the total number of plates/males tested [Mating potency %= (# plates/males with cross progeny/total # of plates/males tested) x 100]. For statistical analysis, we performed a t-test to calculate the significant difference of mating potency between *wild type* worms (*him-5*) and mutant worms (in a *him-5* background).

For backing response after nose touch, we placed one virgin young adult male worm in a clean bacterial lawn and gently touched its nose 10 times with an eyelash, waiting 10 seconds between each touch. The number of times the worm showed response to nose touch by backward movement was scored and the backing response was calculated [Backing response= (# of times the worm backed up after nose touch/10 nose touches) x 100]. For statistical analysis, we performed a t-test to calculate the significant difference of backing response between *wild type* worms (*him-5*) and mutant worms (in a *him-5* background).

### RnB synapses imaging

To visualize the presynaptic pattern of RnB neurons, z-stack images of young adult males containing *Ppkd-2::GFP* and *Ppkd-2::mCherry::RAB-3* reporters (*bxIs30*) were acquired using a Leica SP5 confocal microscope. The z-stack images were analyzed one by one from ventral to dorsal and compressed by looking at the synaptic puncta located in the preanal ganglion. To quantify the protein levels of *mCherry::RAB-3*, we performed a fluorescent densitometry analysis of compressed z-stacks images acquired by using a Zeiss Axioimager Z1 Apotome with a 63x/1.4 objective. For this analysis, we used the same exposure time for all control and mutant samples. The relative fluorescence values were measured by dividing *mCherry* densitometry (corresponding to synapses) by *GFP* densitometry (corresponding to axon terminals). In this way, we corrected for missing synapses that are a product of defects in axonal migration. For statistical analysis, we performed a t-test to calculate the significant difference between control worms and mutant worms.

To visualize the synapses between RnB ray sensory neurons and EF interneurons, we used the iBLINC trans-synaptic biotin transfer system. For the presynaptic RnB sensory neurons labeling, we expressed the biotin ligase with *nrx-1* fusion protein (*BirA::nrx-1*) driven by the *pkd-2/*Polycystin-2 promoter. For the postsynaptic EF interneurons labeling, we expressed the biotinylated acceptor peptide with *nlg-1*/neuroligin fusion protein (*AP::nlg-1*) driven by the *nlg-1*/neuroligin promoter. We co-expressed these pre-and postsynaptic fusion proteins with *streptavidin::RFP* fusion protein driven by the *unc-122* coelomocytes promoter. To quantify RnBs>EFs biotinylated synapses, we performed a fluorescent densitometry analysis of compressed *z*-stack images acquired from mutant and *wild type* worms by using a Zeiss Axioimager Z1 Apotome with a 63x/1.4 objective. Only those worms completely oriented in a dorsoventral position were imaged in order to obtained a clear view of the synaptic ring located in the preanal ganglion. The relative fluorescent (a.u.) values were the measurement of the *RFP* densitometry subtracted by the background (noise). The measured area consists of a circle with a 5 μm radio. To compare the difference in synaptic densities between mutant and *wild type* worms, we divided the relative fluorescence (a.u.) value of mutants by the average of the relative fluorescence (a.u.) of *wild type* worms measured on the same experimental day. For statistical analysis, we performed a *t*-test to calculate the significant difference between *wild type* worms and mutant worms.

### Data Statement

All strains and reagents are available upon request. Figure S1 gives analysis of male potency, response defects and general backing response in the *hst-3.1/*HS 3-*O*-sulfotransferase. Figure S2 provides additional genetic data pertaining to the genetic interactions between *hst-3.1/HS* 3-*O* sulfotransferase, *pkd-2/*polycystin-2, *lov-1/*polycystin-1 and *klp-6/*kinesin. Figure S3 shows expression data of the *hst-3.1/*HS 3-*O*-sulfotransferase transcriptional reporter and *nlg-1*/neuroligin transcriptional GFP fusion. Figure S4 provides additional genetic data pertaining to the role of HSME and HSPG in the axon guidance of B-type ray neurons. Figure S5 provides additional information about the visualization of the B-type ray neuron synapses with the EF interneurons. Figure S6 provides additional genetic data pertaining to the quantification of *mCherry::RAB-3* in the HSME and HSPG mutants. Table S1 provides a complete list of strains created for this study. Tables S2 and S3 provides a complete list of transgenic strains, and the respective constructs, created for this study. File S1-S3 comprises all data used to create figures.

## References

Ackley, B. D., S. H. Kang, J. R. Crew, C. Sue, Y. Jin et al., 2003 The Basement Membrane Components Nidogen and Type XVIII Collagen Regulate Organization of Neuromuscular Junctions in Caenorhabditis elegans. J Neurosci 23:3577–3587.

Allen, N. J., M. L. Bennett, L. C. Foo, G. X. Wang, C. Chakraborty et al., 2012 Astrocyte glypicans 4 and 6 promote formation of excitatory synapses via GluA1 AMPA receptors. Nature 486:410–414.

Altun-Gultekin, Z., Y. Andachi, E. L. Tsalik, D. Pilgrim, Y. Kohara et al., 2001 A regulatory cascade of three homeobox genes, ceh-10, ttx-3 and ceh-23, controls cell fate specification of a defined interneuron class in C. elegans. Development 128: 1951–1969.

Attreed, M., K. Saied-Santiago and H. E. BÜlow, 2016 Conservation of anatomically restricted glycosaminoglycan structures in divergent nematode species.Glycobiology 26: 862–870.

Barr, M. M., J. DeModena, D. Braun, C. Q. Nguyen, D. H. Hall et al., 2001 The Caenorhabditis elegans autosomal dominant polycystic kidney disease gene homologs lov-1 and pkd-2 act in the same pathway. Curr Biol 11: 1341–1346.

Barr, M. M., and P. W. Sternberg, 1999 A polycystic kidney-disease gene homologue required for male mating behaviour in C. elegans. Nature 401: 386–389.

Barrios, A., S. Nurrish and S. W. Emmons, 2008 Sensory regulation of C. elegans male mate-searching behavior. Curr Biol 18: 1865–1871.

Bennett, K. L., J. Bradshaw, T. Youngman, J. Rodgers, B. Greenfield et al., 1997 Deleted in colorectal carcinoma (DCC) binds heparin via its fifth fibronectin type III domain. J Biol Chem 272: 26940–26946.

Bernfield, M., M. Gotte, P. W. Park, O. Reizes, M. L. Fitzgerald et al., 1999 Functions of cell surface heparan sulfate proteoglycans. Annu Rev Biochem 68: 729–777.

Blanchette, C.R., P. N. Perrat,A. Thackeray and C. Y. Benard, 2015 Glypican Is a Modulator of Netrin-Mediated Axon Guidance. PLoS Biol 13: e1002183.

Bose, C. M., D. Qiu, A. Bergamaschi, B. Gravante, M. Bossi et al., 2000 Agrin controls synaptic differentiation in hippocampal neurons. J Neurosci 20: 9086–9095.

Brenner, S., 1974 The genetics of Caenorhabditis elegans. Genetics 77: 71–94.

Bulow, H. E., K. L. Berry, L. H. Topper, E. Peles and O. Hobert, 2002 Heparan sulfate proteoglycan-dependent induction of axon branching and axon misrouting by the Kallmann syndrome gene kal-1. Proc Natl Acad Sci U S A 99: 6346–6351.

BÜlow, H. E., and O. Hobert, 2004 Differential sulfations and epimerization define heparan sulfate specificity in nervous system development. Neuron 41: 723–736.

BÜlow, H. E., and O. Hobert, 2006 The molecular diversity of glycosaminoglycans shapes animal development. Annu Rev Cell Dev Biol 22: 375–407.

BÜlow, H. E., N. Tjoe, R. A. Townley, D. Didiano, T. H. van Kuppevelt et al., 2008 Extracellular sugar modifications provide instructive and cell-specific information for axon-guidance choices. Curr Biol 18: 1978–1985.

Dani, N.M., Nahm, S. Lee and K Broadie, 2012 A targeted glycan-related gene screen reveals heparan sulfate proteoglycan sulfation regulates WNT and BMP trans-synaptic signaling. PLoS Genet 8: e1003031.

de Wit, J., M. L. O’Sullivan, J. N. Savas, G. Condomitti, M. C. Caccese et al., 2013 Unbiased discovery of glypican as a receptor for LRRTM4 in regulating excitatory synapse development. Neuron 79: 696–711.

Desbois, M., S. J. Cook, S. W. Emmons and H. E. BÜlow, 2015 Directional Trans-Synaptic Labeling of Specific Neuronal Connections in Live Animals. Genetics 200: 697–705.

Diaz-Balzac, C. A., M. I. Lazaro-Pena, G. A. Ramos-Ortiz and H. E. BÜlow, 2015 The Adhesion Molecule KAL-1/anosmin-1 Regulates Neurite Branching through a SAX-7/L1CAM-EGL-15/FGFR Receptor Complex. Cell Rep 11: 1377–1384.

Diaz-Balzac, C. A., M. I. Lazaro-Pena, E. Tecle, N. Gomez and H. E. BÜlow, 2014 Complex cooperative functions of heparan sulfate proteoglycans shape nervous system development in Caenorhabditis elegans. G3 (Bethesda) 4: 1859–1870.

Ethell, I. M., F. Irie, M. S. Kalo, J. R. Couchman, E. B. Pasquale et al., 2001 EphB/syndecan-2 signaling in dendritic spine morphogenesis. Neuron 31: 1001–1013.

Ferreira, A., 1999 Abnormal synapse formation in agrin-depleted hippocampal neurons. J Cell Sci 112 (Pt 24): 4729–4738.

Gendrel, M., E. G. Atlas and O. Hobert, 2016 A cellular and regulatory map of the GABAergic nervous system of C. elegans. Elife 5.

Gilleard, J. S., J. D. Barry and I. L. Johnstone, 1997 cis regulatory requirements for hypodermal cell-specific expression of the Caenorhabditis elegans cuticle collagen gene dpy-7. Mol Cell Biol 17: 2301–2311.

Glass, D. J., D. C. Bowen, T. N. Stitt, C. Radziejewski, J. Bruno et al., 1996 Agrin acts via a MuSK receptor complex. Cell 85: 513–523.

Graf, E. R., X. Zhang, S. X. Jin, M. W. Linhoff and A. M. Craig, 2004 Neurexins induce differentiation of GABA and glutamate postsynaptic specializations via neuroligins. Cell 119: 1013–1026.

Hart, M.P., and O. Hobert, 2018 Neurexin controls plasticity of a mature, sexually dimorphic neuron. Nature 553: 165–170.

Hudson, M.L., T. Kinnunen, H. N. Cinar and A. D. Chisholm, 2006 C. elegans Kallmann syndrome protein KAL-1 interacts with syndecan and glypican to regulate neuronal cell migrations. Dev Biol 294: 352–365.

Irie, F., H. Badie-Mahdavi and Y. Yamaguchi, 2012 Autism-like socio-communicative deficits and stereotypies in mice lacking heparan sulfate. Proc Natl Acad Sci U S A 109: 5052–5056.

Jarrell, T. A., Y. Wang, A. E. Bloniarz, C. A. Brittin, M. Xu et al., 2012 The connectome of a decision-making neural network. Science 337: 437–444.

Jia, L., and S. W. Emmons, 2006 Genes that control ray sensory neuron axon development in the Caenorhabditis elegans male. Genetics 173: 1241–1258.

Kastenhuber, E., U. Kern, J. L. Bonkowsky, C. B. Chien, W. Driever et al., 2009 Netrin-DCC, Robo-Slit, and heparan sulfate proteoglycans coordinate lateral positioning of longitudinal dopaminergic diencephalospinal axons. J Neurosci 29: 8914–8926.

Ko, J. S., G. Pramanik, J. W. Um, J. S. Shim, D. Lee et al., 2015 PTPsigma functions as a presynaptic receptor for the glypican-4/LRRTM4 complex and is essential for excitatory synaptic transmission. Proc Natl Acad Sci U S A 112: 1874–1879.

Koo, P. K., X. Bian, A. L. Sherlekar, M. R. Bunkers and R. Lints, 2011 The robustness of Caenorhabditis elegans male mating behavior depends on the distributed properties of ray sensory neurons and their output through core and male-specific targets. J Neurosci 31: 7497–7510.

Ksiazek, I., C. Burkhardt, S. Lin, R. Seddik, M. Maj et al., 2007 Synapse loss in cortex of agrin-deficient mice after genetic rescue of perinatal death. J Neurosci 27: 7183–71951.

Lee, R. Y., E. R. Sawin, M. Chalfie, H. R. Horvitz and L. Avery, 1999 EAT-4, a homolog of a mammalian sodium-dependent inorganic phosphate cotransporter, is necessary for glutamatergic neurotransmission in caenorhabditis elegans. J Neurosci 19: 159–167.

Lin, Y.L., Y. T. Lei, C. J. Hong and Y. P. Hsueh, 2007 Syndecan-2 induces filopodia and dendritic spine formation via the neurofibromin-PKA-Ena/VASP pathway. J Cell Biol 177: 829–841.

Lindahl, U., and J. P. Li, 2009 Interactions between heparan sulfate and proteins-design and functional implications. Int Rev Cell Mol Biol 276: 105–159.

Lints, R., and S. W. Emmons, 1999 Patterning of dopaminergic neurotransmitter identity among Caenorhabditis elegans ray sensory neurons by a TGFbeta family signaling pathway and a Hox gene. Development 126: 5819–5831.

Liu, K. S., and P. W. Sternberg, 1995 Sensory regulation of male mating behavior in Caenorhabditis elegans. Neuron 14: 79–89.

Matsumoto, Y., F. Irie, M. Inatani, M. Tessier-Lavigne and Y. Yamaguchi, 2007 Netrin-1/DCC signaling in commissural axon guidance requires cell-autonomous expression of heparan sulfate. J Neurosci 27: 4342–4350.

Moon, A. F., Y. Xu, S. M. Woody, J. M. Krahn, R. J. Linhardt et al., 2012 Dissecting the substrate recognition of 3-O-sulfotransferase for the biosynthesis of anticoagulant heparin. Proc Natl Acad Sci U S A 109: 5265–5270.

Myers, J. P., M. Santiago-Medina and T. M. Gomez, 2011 Regulation of axonal outgrowth and pathfinding by integrin-ECM interactions. Dev Neurobiol 71: 901–923.

Nguyen, M. U., J. Kwong, J. Chang, V. G. Gillet, R. M. Lee et al., 2016 The Extracellular and Cytoplasmic Domains of Syndecan Cooperate Postsynaptically to Promote Synapse Growth at the Drosophila Neuromuscular Junction. PLoS One 11: e0151621.

Okkema, P. G., S. W. Harrison, V. Plunger, A. Aryana and A. Fire, 1993 Sequence requirements for myosin gene expression and regulation in Caenorhabditis elegans. Genetics 135: 385–404.

Peden, E. M., and M. M. Barr, 2005 The KLP-6 kinesin is required for male mating behaviors and polycystin localization in Caenorhabditis elegans. Curr Biol 15: 394–404.

Pedersen, M. E., G. Snieckute, K. Kagias, C. Nehammer, H. A. Multhaupt et al., 2013 An epidermal microRNA regulates neuronal migration through control of the cellular glycosylation state. Science 341: 1404–1408.

Porcionatto, M. A., 2006 The extracellular matrix provides directional cues for neuronal migration during cerebellar development. Braz J Med Biol Res 39: 313–320.

Poulain, F. E., and H. J. Yost, 2015 Heparan sulfate proteoglycans: a sugar code for vertebrate development? Development 142: 3456–3467.

Pratt, T., C. D. Conway, N. M. Tian, D. J. Price and J. O. Mason, 2006 Heparan sulphation patterns generated by specific heparan sulfotransferase enzymes direct distinct aspects of retinal axon guidance at the optic chiasm. J Neurosci 26: 6911–6923.

Saied-Santiago, K., R. A. Townley, J. D. Attonito, D. S. da Cunha, C. A. Diaz-Balzac et al., 2017 Coordination of Heparan Sulfate Proteoglycans with Wnt Signaling To Control Cellular Migrations and Positioning in Caenorhabditis elegans. Genetics 206: 1951–1967.

Scheiffele, P., J. Fan, J. Choih, R. Fetter and T. Serafini, 2000 Neuroligin expressed in nonneuronal cells triggers presynaptic Development in contacting axons. Cell 101: 657–669.

Sherlekar, A. L., 2015 THE NEURAL AND MOLECULAR MECHANISMS REGULATING MALE LOCOMOTION DURING Caenorhabditis elegans MATING BEHAVIOR, pp. in Biology. Texas A&M University.

Sherlekar, A. L., A. Janssen, M. S. Siehr, P. K. Koo, L. Caflisch et al., 2013 The C. elegans male exercises directional control during mating through cholinergic regulation of sex-shared command interneurons. PLoS One 8: e60597.

Sperry, R. W., 1963 Chemoaffinity in the Orderly Growth of Nerve Fiber Patterns and Connections. Proc Natl Acad Sci U S A 50: 703–710.

SÜdhof, T. C., 2008 Neuroligins and neurexins link synaptic function to cognitive disease. Nature 455: 903–911.

Sulston, J. E., D. G. Albertson and J. N. Thomson, 1980 The Caenorhabditis elegans male: postembryonic development of nongonadal structures. Dev Biol 78: 542–576.

Sze, J. Y., M. Victor, C. Loer, Y. Shi and G. Ruvkun, 2000 Food and metabolic signalling defects in a Caenorhabditis elegans serotonin-synthesis mutant. Nature 403: 560–564.

Tecle, E., C. A. Diaz-Balzac and H. E. Bülow, 2013 Distinct 3-O-sulfated heparan sulfate modification patterns are required for kal-1-dependent neurite branching in a context-dependent manner in Caenorhabditis elegans. G3 (Bethesda) 3: 541–552.

Zimmermann, D. R., and M. T. Dours-Zimmermann, 2008 Extracellular matrix of the central nervous system: from neglect to challenge. Histochem Cell Biol 130: 635–653.

